# A Scalable Touchscreen-Based Spatial Working Memory Task for Cross-Species Research

**DOI:** 10.64898/2026.02.24.707827

**Authors:** Casey R. Vanderlip, Shelby R. Dunn, Payton A. Asch, Craig E.L. Stark, Courtney Glavis-Boom

## Abstract

The common marmoset is rapidly emerging as a powerful nonhuman primate model in neuroscience, yet the development of scalable, mechanistically informative cognitive paradigms has lagged behind advances in neural recording and genetic tools. Here, we introduce and validate a touchscreen-based spatial working memory task designed for direct cross-species translation between marmosets and humans. The paradigm independently manipulates retention delay and spatial separation between test choice stimuli, enabling parametric control over maintenance and interference demands within a single framework. Twelve marmosets and seventy-one human participants performed a Delayed Non-Match-to-Position task in which memory delay (1, 5, 10 seconds) and angular separation between target and distractor locations were systematically varied. Across species, accuracy declined as delay increased and as spatial separation decreased, demonstrating robust sensitivity to both maintenance demands and similarity-based interference. Critically, delay and separation interacted in both species, indicating that these had additive effects. Choice latency analyses further supported interpretation of performance, with slower responses on incorrect trials in both groups. Together, these findings establish a scalable and translationally aligned spatial working memory paradigm that captures interacting maintenance and interference processes. This task provides a powerful platform for circuit-level investigation and offers a sensitive cognitive assay for future studies of aging, neurodegenerative disease, and therapeutic intervention in the marmoset model.

## Introduction

Nonhuman primate models are essential for understanding the neural mechanisms that support higher-order cognition and for translating basic neuroscience findings to humans (Izpisua Belmonte et al., 2015). In this context, the common marmoset (*Callithrix jacchus*) has emerged as an increasingly valuable model species. Marmosets offer several practical advantages over larger primates, including small body size, ease of handling, and relatively short lifespans, enabling experimental and longitudinal designs that are difficult to achieve in other nonhuman primates (Perez-Cruz & Rodriguez-Callejas, 2023; Tardif, Mansfield, Ratnam, Ross, & Ziegler, 2011). Importantly, these benefits are paired with key features of primate brain organization, including a differentiated prefrontal cortex and conserved frontoparietal networks that support executive functions, such as working memory (Glavis-Bloom, Vanderlip, & Reynolds, 2022; Vanderlip, Asch, Reynolds, & Glavis-Bloom, 2023; Watakabe et al., 2023).

Despite rapid growth in marmoset-based research, progress has been constrained by a limited number of cognitive tasks optimized for this species and suitable for translational research (Chen, Menegas, Zhang, & Feng, 2025). Many commonly used behavioral paradigms were developed for rodents or larger primates and do not readily scale to marmosets. Such limitations restrict the ability to probe cognitive mechanisms, assess parametric effects on behavior, or link performance to underlying neural circuitry. Therefore, there is a pressing need for cognitive tasks that are efficient, flexible, and capable of systematically modulating cognitive demand.

Working memory is a core component of executive function, enabling the temporary maintenance and manipulation of information to guide goal-directed behavior (Baddeley, 2003; Cohen et al., 1997; Miller, Li, & Desimone, 1991). In primates, working memory depends critically on prefrontal and frontoparietal circuitry and can be challenged along multiple dimensions, including the duration over which information must be maintained and the degree of interference between competing representations (Goldman-Rakic, 1996). Parametric manipulation of these dimensions provides a powerful approach for taxing working memory processes and isolating circuit-level contributions to performance. Tasks that support such manipulations within a single framework are particularly well-suited for translational and systems neuroscience.

Working memory tasks have been used in marmosets using delayed response and delayed non-match paradigms, demonstrating that marmosets can maintain information across short delays and engage prefrontal circuitry during memory tasks (Miles, 1957; Spinelli et al., 2004; Wong, Selvanayagam, Johnston, & Everling, 2023; Yamazaki, Saiki, Inada, Watanabe, & Iriki, 2016). However, existing paradigms do not manipulate both delay and interference in the same paradigm, limiting mechanistic interpretation of whether performance costs reflect weakened maintenance versus increased competition at retrieval. A task that independently manipulates both dimensions offers stronger leverage for circuit-level studies and improves sensitivity to subtle impairment. In addition, working memory tasks in marmosets have not been implemented in humans using matched task parameters, limiting direct cross-species comparison.

Here, we introduce a touchscreen-based spatial working memory task developed for use in marmosets and designed for direct translatability in humans. Conceptually, the paradigm draws inspiration from spatial foraging and navigation tasks such as the radial “cheeseboard” task and the Barnes maze, which assess memory for location within an allocentric spatial array (Barnes, 1979; Gilbert, Kesner, & Lee, 2001; Kesner, 2007). Our implementation translates these spatial discrimination principles into a controlled touchscreen format that allows independent manipulation of memory delay and the spatial proximity of choice stimuli. This flexibility makes it possible to tune task difficulty, generate rich behavioral distributions, and probe working memory function across a range of cognitive demands rather than at a single operating point. Together, this work provides a flexible and translational spatial working memory paradigm that expands the cognitive toolkit available for marmoset research, and lends itself to direct human translation.

## Methods

### Subjects

#### Marmosets

Twelve common marmosets (*Callithrix jacchus*; 6 males, 6 females) between the ages of 6.1 and 14 years (mean = 9.46 years; SD = 2.36 years) participated in the study. This age range spans mid-adulthood through geriatric stages in marmosets, roughly corresponding to approximately 30 to 80+ years in humans based on lifespan scaling estimates. All marmosets had extensive prior experience with touchscreen-based cognitive testing in their home cages (Vanderlip et al., 2023). Animals were either singly or pair-housed in cages with environmental enrichment (hammocks and manzanita branches), as well as visual and auditory access to other marmosets. During testing, pair-housed animals were temporarily separated by a solid divider in the home cage. All experimental procedures were approved by the Salk Institute Animal Care and Use Committee and followed the guidelines of the NIH Guide for the Care and Use of Laboratory Animals.

#### Humans

Seventy-one people (12 males, 59 females) between the ages of 18.72 and 40.45 years (mean = 22.46 years; SD = 4.71 years) participated in the study. Participants were recruited from the University of California, Irvine undergraduate subject pool (SONA) and through word-of-mouth advertisement within the university community. Following data-quality filtering to exclude subjects with performance patterns consistent with low task engagement, sixty subjects (9 males, 51 females) within the same age range (mean = 22.69, SD = 5.08 years) remained. All experimental procedures were approved by the University of California, Irvine Institutional Review Board.

### Equipment

#### Marmosets

Marmoset home cages were custom-built to include a testing chamber located in the top corner of each cage, where a 10.4-inch infrared touchscreen apparatus (Lafayette Instrument Company, Lafayette, IN) was mounted. Access to this testing chamber was restricted outside of cognitive testing sessions, during which the marmosets could freely enter and exit the chamber through a small doorway. Liquid rewards (e.g., apple juice) for correct responses were dispensed into a small basin mounted beneath the touchscreen via a peristaltic pump. Stimulus locations on the screen were defined by task-specific cutouts in a plastic overlay placed in front of the display, providing a consistent visual frame for response options. All tasks were programmed and administered using Animal Behavior Environment Test (ABET) Cognition software (Lafayette Instrument Company, Lafayette, IN). The software recorded millisecond resolution data including response accuracy, location of stimuli and touches on the screen.

#### Humans

The human version of the DNMP task was implemented using jsPsych (v6.3.1) and administered online via JATOS. Participants completed the task in full screen mode on their personal computers.

### Cognitive Testing

Marmosets and humans were tested on a Delayed Non-Match-to-Position (DNMP) task to assess spatial working memory (Figure 1). The marmoset and human versions of the task were similarly structured and parameterized to enable cross-species comparisons and promote future translational research.

**Figure 1.**
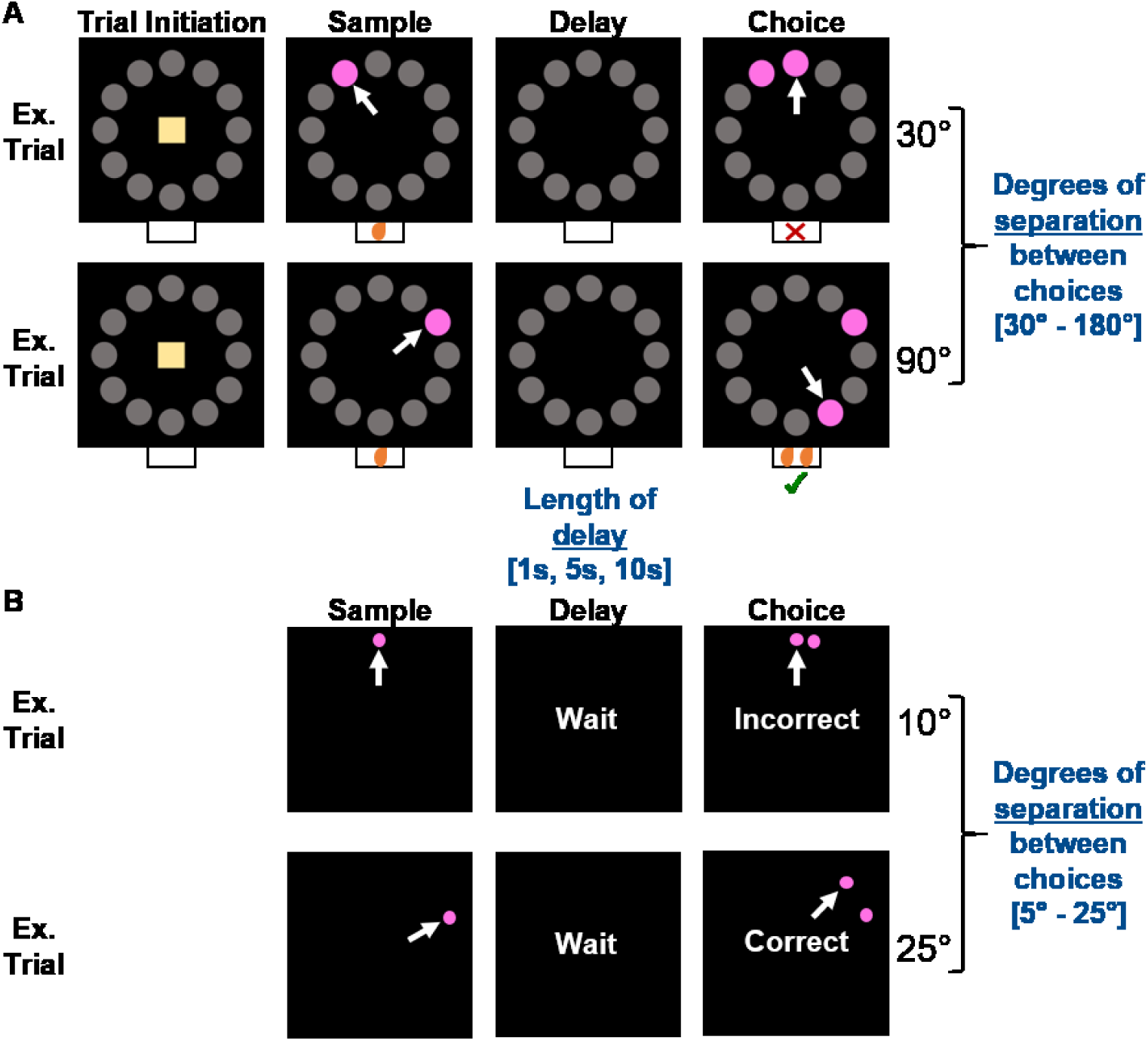
Delayed Nonmatch-to-Position (DNMP) task schematics. A) Examples of two trial progressions for marmosets and B) humans. Trial difficulty was modulated by delay (1s, 5s, 10s) and the radial separation between choices. Marmoset choices were separated by 30° to 180° in 30° increments. Human choices were separated by 5° to 25° in 5° increments.

#### Marmosets

A black plastic “mask” placed in front of the touch screen had cutouts delineating the potential locations in which stimuli could appear on the screen. The cutouts were arranged in a circular pattern, with 12 locations spaced equidistant in typical clock face locations, and an additional cutout in the center of the screen for the trial initiation stimulus to be presented. Each trial was initiated by the subject selecting a trial initiation stimulus in the center of a black screen. Then, one of the 12 locations was illuminated by a solid pink stimulus. Selection of this sample stimulus resulted in a blank screen, delivery of a small liquid reward, and the start of a 1, 5, or 10s retention delay. Following the delay, the original location and a novel location were illuminated by identical solid pink stimuli. Selection of the stimulus in the novel location resulted in a correct response, delivery of a liquid reward, and the start of a 5s inter-trial interval. Incorrect selections (choosing the original location) or omissions (no response within 12s) resulted in no reward being dispensed and triggered a 5s timeout in addition to the 5s inter-trial interval.

Given that the stimuli were visually identical, marmosets could rely only on spatial information to determine the correct response. Using a radial array for potential stimulus locations allowed us to parametrically manipulate angular separation between the two choice stimuli from 30° to 180° in 30° increments. Varying both the length of the retention delay and the spatial proximity provided a controlled way to titrate interference and spatial working memory load, enabling assessment of main effects and interactions between these parameters on task performance.

Marmosets were tested 1 to 3h per day, 3 to 5 days per week, and were not subjected to food or water restriction. Each daily testing session began with a training phase. Trials in the training phase used a 1s delay and larger separations between choice locations (≥ 90°) to minimize interference and facilitate acquisition of the task rule (selection of the novel stimulus). Once marmosets achieved a criterion of 18 correct responses out of 20 consecutive trials, marmosets proceeded to the testing phase. Testing phase trials included all possible separations between choices (30° to 180° in 30° increments) and all delays (1s, 5s, and 10s). Marmosets completed approximately 200 trials of each unique combination of delay and separation, with testing delays presented in blocked order (1s, then 5s, then 10s) and separations presented pseudorandomly. Trial-level data included original and novel locations, separation level, delay duration, response accuracy, and choice latency, defined as the time from choice stimuli onset to selection.

#### Humans

The human version of the DNMP task was closely matched to that of the marmosets. Potential locations for stimuli to appear were presented within a 600 x 600 pixel circular display with 72 equally spaced possible locations, therefore allowing for separations between choice stimuli in 5° increments. Humans were given brief written instructions on the screen prior to the task beginning and completed a total of 50 testing phase trials, with separations of 5° to 25° in 5° increments, and identical delays to the marmosets (1s, 5s, and 10s). The delay and separation for each trial were selected randomly. There was no trial initiation, and feedback following choice selection was delivered via the written word “Incorrect” in red font or “Correct!” in green font on the screen for 1s. Trial-level data included the same variables as for marmosets.

### Statistical methods

Data were analyzed using R Version 4.5.0 (R Core Team, 2023). Trial-level accuracy (correct vs incorrect) was analyzed separately in marmosets and humans using generalized linear mixed-effects models (GLMMs) with a binomial error distribution and using the logit link function. This approach models the probability of a correct response while accounting for repeated trials within subjects. In the primary model, delay was treated as a categorical factor (1s, 5s, 10s), and separation (degrees) was treated as a continuous predictor that was mean-centered and scaled to z-scores to facilitate interpretation. Sum-to-zero contrasts were applied prior to model fitting to enable interpretable Type III tests of main effects and interactions. Fixed effects included delay, separation, and their interaction to test whether the effect of separation depended on delay. To account for within-subject dependence and individual differences, models included a random intercept for each subject and a random subject-specific slope for separation (uncorrelated with the intercept), allowing subjects to differ in overall accuracy and in how strongly separation influenced performance. Models were fit by maximum likelihood using the Laplace approximation.

Post hoc inference focused on three questions. First, differences in accuracy across delay levels were evaluated using estimated marginal means on the response (probability) scale with Tukey-adjusted pairwise comparisons. Second, the overall effect of separation was evaluated using estimated marginal trend methods (emtrends), with a one-sided test of the hypothesis that smaller separation between choices decreases performance (i.e., positive slope for separation). Third, to test whether the effect of separation differed across delay levels, simple slopes of separation were estimated within each delay level and compared using Holm-adjusted pairwise tests to control family-wise error. For interpretability, separation effects were reported as odds ratios per +1 SD increase in separation, with Wald 95% confidence intervals.

To test whether age moderated task effects in marmosets, age was included as a continuous predictor in generalized linear mixed-effects models (binomial logit link) that incorporated interactions among delay, separation, and age (delay × separation × age). Delay was modeled as a categorical factor and separation as a continuous predictor. The models included random intercepts and a random slope for separation by animal, preserving the repeated-measures structure of the trial-level data while allowing animals to differ in baseline performance and sensitivity to separation. Moderation was evaluated using Type III Wald tests and model-based estimated marginal means and trends. Model adequacy was assessed using standard diagnostics including checks for overdispersion, collinearity, convergence warnings, and singular random-effects fits.

Latency was analyzed using linear mixed-effects models. Latencies greater than 12 seconds were excluded. Delay was modeled as a categorical factor, and accuracy was coded as correct versus incorrect. Models included fixed effects of accuracy, delay, and their interaction, with a random intercept of subject to account for repeated measures. Models were estimated using restricted maximum likelihood (REML), and inference was based on Satterthwaite approximations. Type III tests were conducted with planned comparisons that examined differences in latency between correct and incorrect responses at each delay (Holm-adjusted).

### Data curation

Marmoset data were filtered prior to analysis by excluding trials on which the marmoset failed to make a response within 12s of stimulus presentation. This resulted in an average of 6.47% of trials being removed (SD = 3.55%, range: 1.65% to 13.9%).

Human data were filtered prior to analysis by excluding subjects whose mean accuracy on the 1s delay trials was < 60%, consistent with insufficient task engagement. This resulted in 3 males and 8 females being removed.

### Transparency and openness

Data and analysis code is available from the corresponding author upon reasonable request. The design and analyses for this study were not preregistered.

## Results

### Extending delay reduces accuracy in both marmosets and humans

Across species, accuracy decreased as the retention interval (i.e., delay) increased. In marmosets (Figure 2A), the random-slope GLMM (random intercept + random slope for separation) showed a robust main effect of delay (χ²(2) = 482.89, p < 2.2 × 10^−16^). In the categorical-delay model, both longer delays (5s and 10s) were associated with lower accuracy relative to the 1s delay (1s vs 5s: odds ratio = 1.41, p < 0.0001; 1s vs 10s: odds ratio = 1.93, p < 0.0001), and accuracy at 5s was also higher than at 10s (odds ratio = 1.37, p < 0.0001). Estimated marginal means showed accuracy decreasing monotonically across delays (1s delay: p = 0.768 ± 0.00769; 5s delay: p = 0.701 ± 0.00889; 10s delay: p = 0.632 ± 0.00989), with all pairwise delay contrasts significant (Tukey-adjusted p < 0.0001).

**Figure 2.**
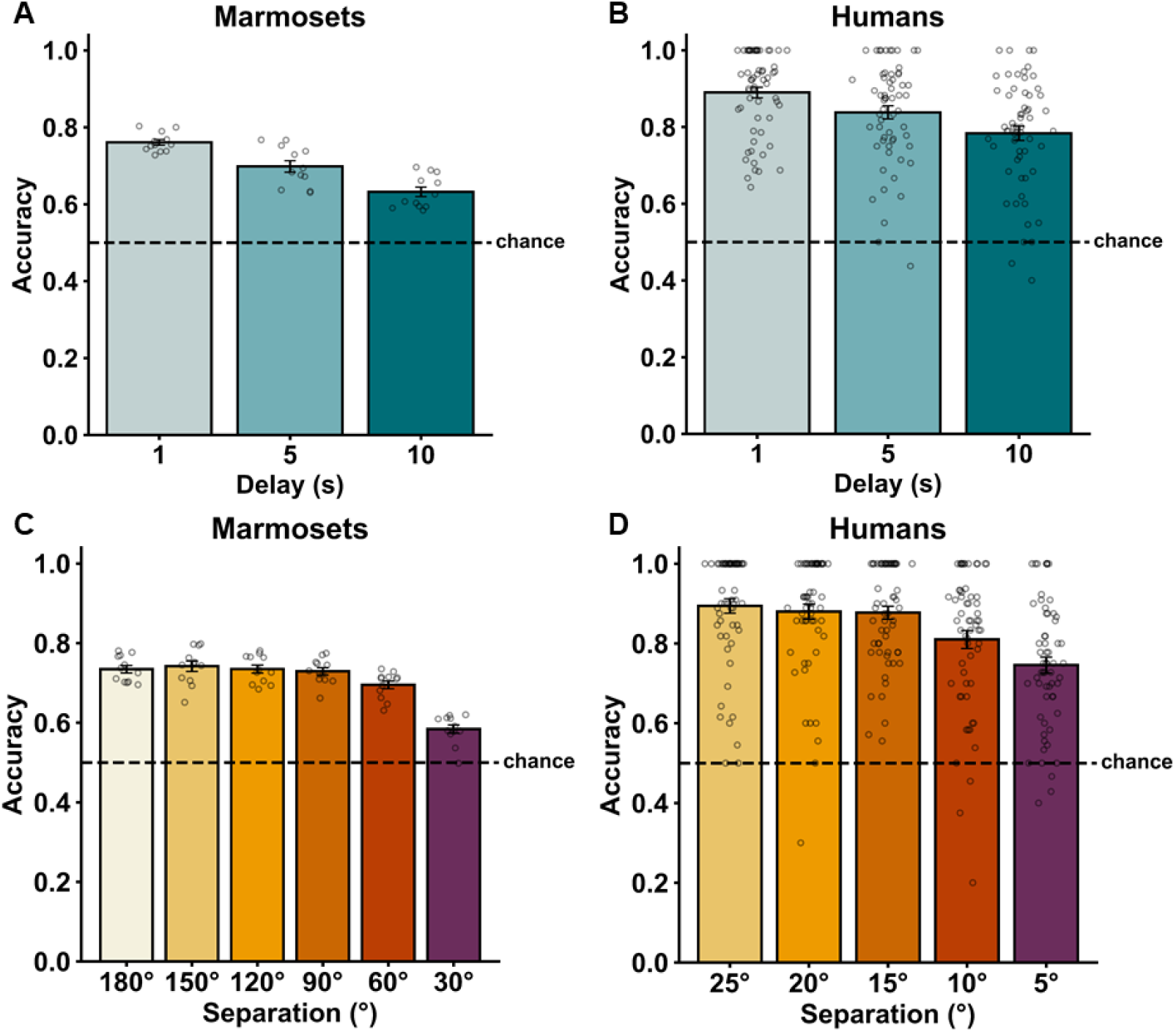
Accuracy decreases as a function of delay and separation in marmosets and humans. A) Accuracy on the Delayed Non-Match-to-Position (DNMP) task significantly decreases as delay increases in marmosets and B) humans. C) Accuracy on the DNMP task significantly decreases as the radial separation between the choices decreases in marmosets and D) humans.

Humans showed the same delay-related decrement in accuracy (Figure 2B). In the categorical-delay GLMM, there was a significant main effect of delay (χ²(2) = 37.64, p = 6.70 × 10^−9^). Relative to performance at the 1s delay, accuracy was significantly lower at 5s (odds ratio = 1.51, p = 0.01) and further reduced at 10s (odds ratio = 2.28, p < 0.0001), with accuracy at 5s also higher than at 10s (odds ratio = 1.51, p = 0.0035), indicating increasing memory demand produced reliable performance costs.

### Less separation between choices reduces accuracy in both marmosets and humans

Accuracy also depended on the amount of spatial separation between the choices. In marmosets, smaller separation was strongly associated with lower accuracy in the GLMM (β = 0.234, SE = 0.0164, z = 14.25, p < 2 × 10^−16^), such that decreasing separation reliably reduced the probability of a correct response (Figure 2C). In humans, the same pattern was observed (Figure 2D). Specifically, a one standard deviation decrease in separation was associated with a 35% reduction in the odds of a correct response (β = 0.426, SE = 0.0695, z = 6.14, p = 8.34 × 10^−10^), indicating that closer spatial proximity impaired performance in both species.

### Spatial interference increases with delay in both marmosets and humans

For both marmosets and humans, delay and separation interacted, indicating that the effect of separation depended on the delay interval. In marmosets, the delay × separation interaction was significant in the categorical-delay GLMM (Figure 3A; χ²(2) = 102.80, p < 2 × 10^-16^). Consistent with this interaction, the separation slope was strongest at the shortest delay and progressively weakened as delay increased. The estimated simple slopes for separation were largest at the 1s delay and smaller at the 5s and 10s delays (1s delay: β = 0.400, SE = 0.0249, odds ratio = 1.49, 95% CI [1.42, 1.57]; 5s delay: β = 0.207, SE = 0.0232, odds ratio = 1.23, 95% CI [1.18, 1.29]; 10s delay: β = 0.0953, SE = 0.0232, odds ratio = 1.10, 95% CI [1.05, 1.15]). These separation slopes differed significantly across delay levels (Holm-adjusted: 1s vs 5s, p < 0.0001; 1s vs 10s, p < 0.0001; 5s vs 10s, p = 0.0001), indicating that closer separations were most detrimental for accuracy at the shortest delay and least detrimental for accuracy at the longest delay. Delay differences were present across separations, with accuracy lowest at small (1 SD below mean) separations, and highest at large (1 SD above mean) separations (all Tukey-adjusted p-values ≤ 0.0003).

**Figure 3.**
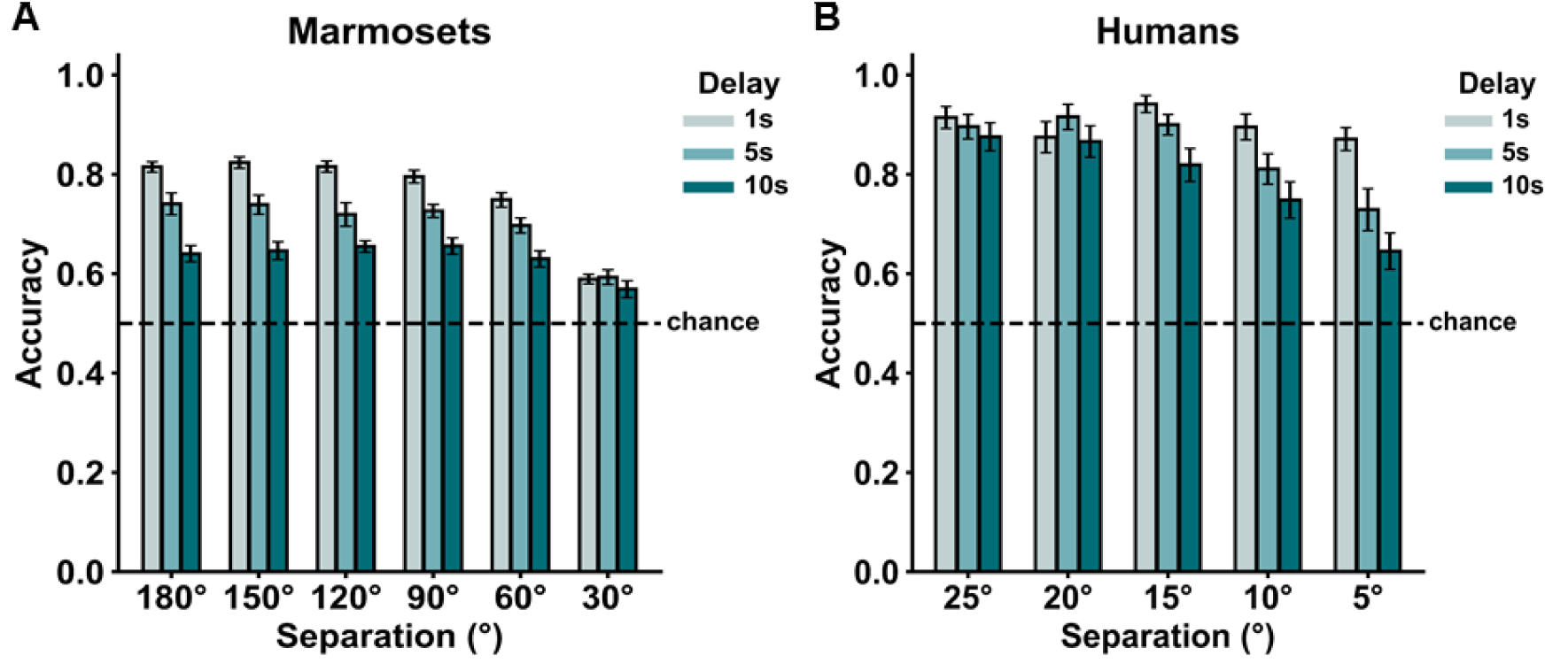
Significant interaction between delay and separation on accuracy. A) Accuracy as a function of delay and separation distance for marmosets and B) humans. In both species, the effect of separation on accuracy depended on the length of the delay. Performance accuracy was highest on trials with a 1s delay and large separation between choices and lowest on trials with a 10s delay and small separation between choices. The longer delays exacerbated the detrimental effects of small spatial separation between choice stimuli, consistent with spatial interference increasingly affecting performance as memory demands increase.

Humans showed a similar interaction between separation and delay, with smaller separations resulting in greater performance costs as delay increased, but this was more modest than in marmosets (Figure 3B). In the categorical-delay GLMM, the delay × separation interaction was marginal overall (χ²(2) = 5.78, p = 0.055), indicating a weaker moderation effect than observed in marmosets. Simple-slope analyses clarified the pattern: the effect of separation was smallest at the 1s delay (β = 0.232, SE = 0.114, odds ratio = 1.26) and stronger at the 5s and 10s delays (5s delay: β = 0.508, SE = 0.104, odds ratio = 1.66, 95% CI [1.36, 2.04]; 10s delay: β = 0.540, SE = 0.094, odds ratio = 1.72, 95% CI [1.43, 2.06]). In summary, a 1 SD decrease in separation between choice stimuli was associated with progressively more reduction in the odds of a correct response at longer delays.

### Age did not moderate task performance in marmosets

Since the marmoset subjects spanned from adult to geriatric, we tested whether age moderated the effects of delay and separation on accuracy using mixed-effects logistic regression models with random intercepts and a random slope for separation by animal. Controlling for age, longer delays were still associated with lower accuracy (Type III: χ²(2) = 481.24, p < 2×10^-16^), and larger separation between choice stimuli was associated with higher accuracy (χ²(1) = 227.19, p < 2×10^-16^). The delay × separation interaction also remained when controlling for age (χ²(2) = 102.36, p < 2×10^-16^), indicating that the effect of separation was strongest at shorter delays, regardless of age.

When testing directly for effects of age, there was no significant main effect of age (χ²(1) = 0.87, p = 0.35), nor was there evidence that age moderated separation effects (age × separation: χ²(1) = 0.90, p = 0.34) or the delay × separation interaction (age × delay × separation: χ²(2) = 3.47, p = 0.18). A small age × delay interaction was detected (χ²(2) = 7.17, p = 0.028). However, this effect was minimal in magnitude (on the order of approximately one to three percentage points) and not visually appreciable. Overall, the magnitude and form of delay-and separation-related performance accuracy changes were largely consistent across the age range tested in marmosets. We did not perform age analyses in humans given the restricted age range.

### Accuracy and delay modulate choice latency in marmosets and humans

In marmosets, there was a significant main effect of trial accuracy on choice latency such that incorrect responses were slower than correct responses (Figure 4A; χ²(1) = 235.82, p < 0.001). There was also a main effect of delay (χ²(2) = 5561.50, p < 0.001). Overall, latency on 1s delay trials was significantly slower than on both 5s and 10s delay trials (both p < 0.001), and latency on 5s delay trials was significantly faster than on 10s delay trials (p < 0.001). Critically, the accuracy × delay interaction was significant, indicating that the magnitude of the correct vs incorrect latency difference varied across delays (χ²(2) = 8.34, p = 0.015). Pairwise comparisons confirmed that incorrect responses were significantly slower than correct responses on trials of all delay lengths (1s: β = −0.301, p < 0.001; 5s: β = −0.317, p < 0.001; 10s: β = −0.202, p < 0.001).

**Figure 4.**
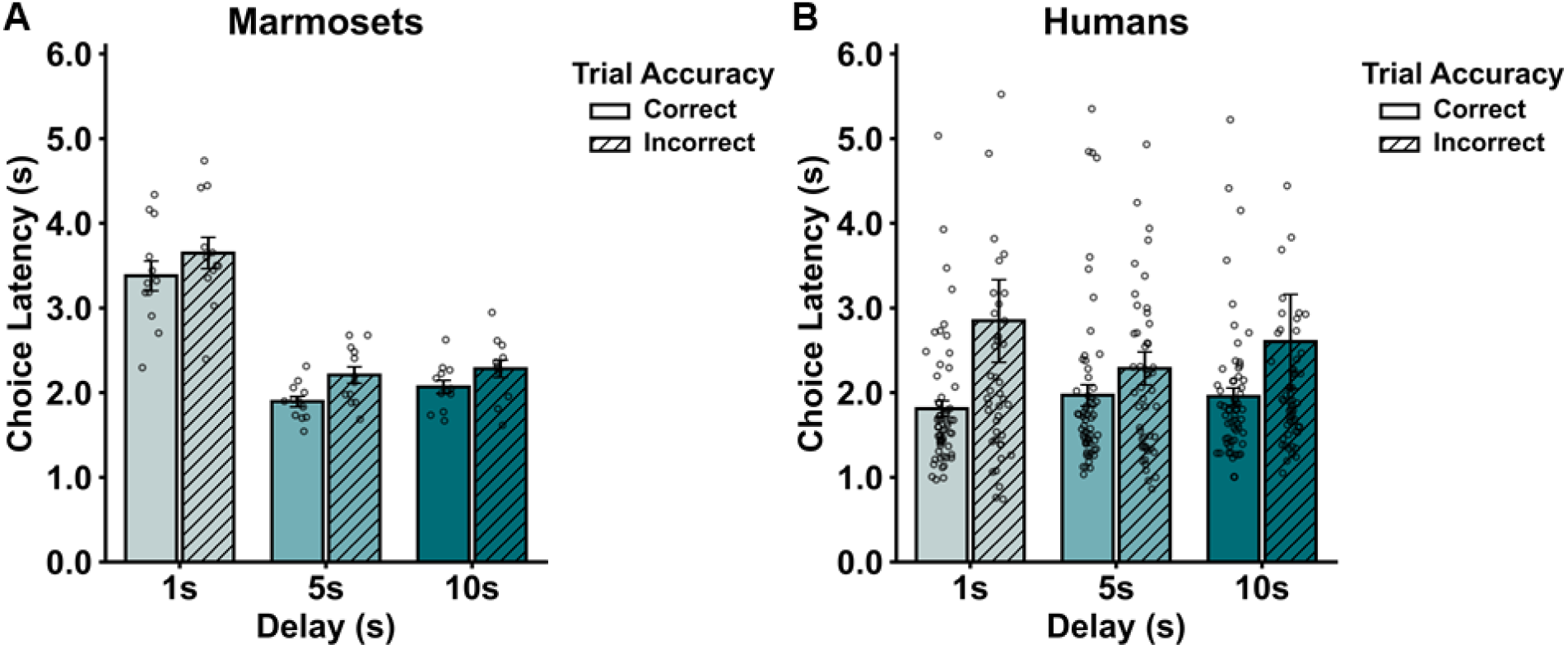
Significant interaction between delay and trial accuracy on choice latency. A) Choice latency as a function of delay and correct vs incorrect trials for marmosets and B) humans. In both species, incorrect response choice latencies were longer than correct choice latencies, and the magnitude of this difference depended upon delay.

In humans, there was also a significant main effect of trial accuracy on choice latency such that incorrect responses were slower than correct responses (Figure 4B; χ²(1) = 32.67, p < 0.001). Unlike marmosets, there was no significant main effect of delay (χ²(2) = 2.39, p = 0.303). However, like marmosets, the accuracy × delay interaction was significant, indicating that the effect of correct vs incorrect choice on latency depended on delay (χ²(2) = 14.47, p < 0.001). Pairwise comparisons revealed that incorrect responses were significantly slower than correct responses on trials with a 1s and 5s delay, but not on trials with a 10s delay (1s delay: β = −0.579, p < 0.001; 5s delay: β = −0.282, p = 0.002; 10s delay: β = −0.077, p = 0.332).

## Discussion

In this study, we developed and validated a touchscreen-based spatial working memory task explicitly for translational use in marmosets and humans. By independently manipulating delay and spatial separation between choice stimuli, the paradigm provides parametric control over two core dimensions of working memory demand: maintenance and interference. Across species, performance declined systematically as delay increased and as spatial separation decreased. Moreover, delay and separation interacted, demonstrating that maintenance-related degradation and similarity-based competition jointly impact working memory performance. Together, these findings establish the task as a scalable, mechanistically informative, and translationally aligned platform for investigating spatial working memory across species.

### Parametric control of working memory demand

Compared with prior delayed response and delayed non-match paradigms used in marmosets and other primates, a key feature of the present task is the explicit manipulation of spatial separation between competing choice stimuli (Spinelli et al., 2004; Wong et al., 2023; Yamazaki et al., 2016). Many existing working memory tasks vary delay but hold similarity constant, limiting the ability to systematically tax retrieval competition. In the present design, angular separation between the target and foil is varied parametrically, modulating the degree of similarity at test and thereby scaling task difficulty. Smaller separations increase the similarity between the encoded location and the alternative, placing greater demands on the precision of the maintained spatial representation. This manipulation allows interference at retrieval to be experimentally titrated within a single framework, enabling assessment of how representational fidelity interacts with delay. Such graded control over retrieval difficulty may enhance sensitivity to subtle changes in working memory precision that are not captured by delay manipulations alone. This additional degree of control may therefore enhance sensitivity to subtle disease-related changes that would not be captured by delay manipulations alone (Gazzaley, Cooney, Rissman, & D’Esposito, 2005; Stopford, Thompson, Neary, Richardson, & Snowden, 2012a; Vanderlip, Stark, & for the Alzheimer’s Disease Neuroimaging Initiative, 2024).

In alignment with prior work, increasing the delay interval reliably reduced accuracy in both marmosets and humans, consistent with progressive weakening of maintained spatial representations over time (Dumitriu et al., 2010; Gallagher & Rapp, 1997; Morrison & Baxter, 2012; Vanderlip et al., 2025). Decreasing angular separation between choice locations similarly impaired performance, reflecting increased spatial interference at retrieval. Because stimuli were visually identical and differed only in spatial position, successful performance depended entirely on maintaining and discriminating spatial representations rather than relying on perceptual cues.

Performance did differ across species, with humans performing at higher accuracy than marmosets across conditions. The human sample was composed of young adults, whereas the marmoset cohort spanned adult to geriatric ages, which may also contribute to baseline performance differences. The human version also included more potential stimulus locations and finer separation increments. Our goal was not to equate absolute task difficulty across species, but to apply parallel manipulations of delay and spatial separation to probe similar cognitive computations across species. Differences in baseline accuracy therefore reflect performance range rather than differences in the structure of the task. Notably, if closer alignment of performance levels were desired, delay intervals could be extended in humans to shift performance away from ceiling and increase sensitivity to separation effects.

The interaction between delay and separation further clarifies how these manipulations operate within each species. In marmosets, the effect of spatial separation was strongest at short delays and weaker at longer delays. In humans, the opposite pattern was observed, with separation effects most evident at the longest delays. One possible explanation for this difference is the location of each species along the performance continuum. In humans, accuracy at short delays neared ceiling levels across many of the separation conditions, limiting the observable impact of interference. As delay increased and performance moved away from ceiling, separation effects became more apparent. In marmosets, performance at long delays approached lower accuracy levels, which may have reduced the measurable contribution of separation under those conditions.

Importantly, both species showed that the impact of spatial separation depended on delay, indicating that maintenance and interference do not operate independently. Rather than equating difficulty across species, the key finding is that manipulating delay and similarity alters performance in systematic and interacting ways in both marmosets and humans. This shared structure supports the conclusion that the task engages comparable working memory computations across species while allowing performance to vary within each species’ operating range.

Such validation is critical for translational neuroscience (Hargreaves, Mattfeld, Stark, & Suzuki, 2012; Vanderlip, Asch, & Glavis-Bloom, 2024; Vanderlip, Dunn, Asch, & Glavis-Bloom, 2026). Nonhuman primate models are often assumed to recapitulate human cognitive processes, yet direct behavioral alignment is rarely demonstrated under identical parametric manipulations. The present findings provide evidence that marmosets and humans exhibit comparable sensitivity to maintenance demands and spatial interference within the same task framework. This alignment strengthens the interpretability of future neural, pharmacological, and genetic studies conducted in the marmoset model.

### Latency as an index of performance

Computerized cognitive assessment enables measurement of behavior beyond accuracy, including trial-level response timing (Glavis-Bloom et al., 2022; Reber, Alvarez, & Squire, 1997). Here, choice latency analyses provided converging evidence that errors in this task reflect degraded or ambiguous memory representations rather than impulsive responding. In both marmosets and humans, incorrect responses were consistently slower than correct responses. This pattern suggests that errors arise when subjects require additional processing time to resolve competition between alternatives, consistent with weakened or overlapping spatial representations. Specific to the marmosets, the choice latencies on trials with a 1s delay were longer than those on trials with a 5s or 10s delay. We attribute this to the fact that marmosets received a reward for selecting the sample and likely required more than the 1s delay to consume the reward prior to making a response, thus elongating the response times for those trials. Overall, the availability of trial-level latency measures enhances the paradigm’s value, as response timing can be directly related to neural indices of representational strength, interference, and decision dynamics.

### Implications for aging and Alzheimer’s disease research

The marmoset has gained increasing traction as a model for age-related neurodegenerative disease, including Alzheimer’s disease (Homanics et al., 2024; Huhe et al., 2025; Sukoff Rizzo et al., 2023). Its relatively short lifespan, primate cortical organization, and emerging transgenic lines position it uniquely among nonhuman primates for modeling human neurodegenerative processes. As transgenic marmoset models of Alzheimer’s pathology continue to develop, there is a pressing need for sensitive, scalable cognitive paradigms capable of detecting subtle working memory impairments across disease progression.

Working memory dysfunction emerges early in Alzheimer’s disease and is strongly linked to prefrontal and frontoparietal network integrity (Baddeley, Bressi, Della Sala, Logie, & Spinnler, 1991; Gazzaley et al., 2005; Satler, Belham, Garcia, Tomaz, & Tavares, 2015; Stopford, Thompson, Neary, Richardson, & Snowden, 2012b). The present task is well suited to interrogate these systems. Because it parametrically modulates maintenance and interference demands, the paradigm can detect not only overall performance decline but also shifts in sensitivity to delay or spatial similarity. Such dissociations may be particularly informative in disease contexts. For example, pathological changes that preferentially degrade representational stability might amplify delay-related effects, whereas alterations in inhibitory control or network competition might disproportionately increase interference sensitivity.

Moreover, the task’s scalability and high trial yield make it appropriate for longitudinal designs, enabling within-subject tracking of cognitive change over time. In the context of emerging transgenic models, this paradigm could serve as a standardized behavioral assay for assessing disease onset, progression, and therapeutic intervention effects. Aligning the same task across marmosets and humans also provides a bridge for translating preclinical findings to human cognitive phenotypes.

### A platform for circuit-level and systems neuroscience

Working memory remains central to debates regarding persistent activity, dynamic coding, and distributed network representations (Curtis & D’Esposito, 2003; Curtis & Sprague, 2021; Lundqvist, Herman, & Miller, 2018; Miller, Lundqvist, & Bastos, 2018). Addressing these questions requires tasks that can systematically modulate cognitive demand while remaining compatible with neural recording and manipulation techniques. The present paradigm provides this leverage. Independent control over delay and separation enables targeted interrogation of maintenance-related and interference-related processes. Neural activity can be examined as a function of each dimension, as well as their interaction, providing a powerful framework for linking circuit dynamics to behavior.

The touchscreen-based implementation supports high trial counts and can be readily integrated with electrophysiology, calcium imaging, optogenetic or chemogenetic interventions, and eye tracking (Kangas & Bergman, 2017; Talpos, McTighe, Dias, Saksida, & Bussey, 2010; Vanderlip, Asch, et al., 2024). Such flexibility positions the task as more than a behavioral assay; it is a platform for mechanistic investigation.

## Conclusion

This work establishes a robust, scalable, and translationally aligned spatial working memory paradigm that parametrically manipulates maintenance and interference demands. The task produces convergent behavioral effects in marmosets and humans, supports fine-grained measurement through accuracy and latency, and is well suited for integration with systems-level approaches. As marmosets increasingly serve as models for aging and Alzheimer’s disease, the availability of sensitive and mechanistically grounded cognitive tools becomes essential. The present paradigm provides a powerful foundation for future investigations of working memory circuitry, neurodegeneration, and translational neuroscience.

## Acknowledgments

This research was supported by an AHA-Allen Initiative in Brain Health and Cognitive Impairment award made jointly through the American Heart Association and The Paul G. Allen Frontiers Group: 19PABH134610000AHA, National Institutes of Health grants 1R21AG068967-01, R01AG066683 (CS), P30AG066519 (CS) and T32AG00096 (CS, CV), grants from the Larry L. Hillblom Foundation and the Don and Lorraine Freeberg Foundation, and the Fiona and Sanjay Jha Chair in Neuroscience. We thank Katie Williams for assistance in the care of the marmosets and technical support.

